# Altered Somatosensory Cortical Entrainment in Cerebral Palsy

**DOI:** 10.64898/2026.10.19.760247

**Authors:** Owais A. Khan, Sarah E. Baker, Elizabeth Heinrichs-Graham, Tony W. Wilson, Max J. Kurz

## Abstract

Somatosensory dysfunction contributes substantially to mobility impairments in cerebral palsy (CP), yet their somatosensory cortical responses to rhythmic sustained input remains unstudied. This study examined somatosensory cortical entrainment in 28 ambulatory youth with CP (Gross Motor Function Classification System levels I–III; 21.9±6.3y; 14 unilateral CP) and 34 neurotypical controls (23.4±3.8y). Participants underwent magnetoencephalography while an 8 Hz cutaneous electrical stimulation was applied to the right posterior tibial nerve. Rhythmic neural responses were mapped to the contralateral somatosensory cortices using beamforming, and virtual sensor time series were extracted to quantify neural dynamics. The entrainment response declined more steeply over time in CP compared to controls, whereas the concurrent alpha response strengthened progressively in CP but changed minimally in controls (Group×Time interactions, *p*s<.001). Significant cross-spectral coupling of entrainment×concurrent alpha responses was observed only in CP (*p*<.001) and was most pronounced in those with bilateral CP. Weaker cross-spectral coupling was associated with better functional mobility in controls (*p*=.004) but brain-behavior links in CP were phenotype-specific. Finally, the second harmonic neural response in CP was weaker (*p*<.001) and demonstrated a blunted temporal modulation relative to controls (*p*=.003). These findings implicate disrupted somatosensory cortical entrainment responses as possible neurophysiological features of youth with CP.

---

Cerebral palsy (CP) is conventionally characterized by altered developmental trajectories and lifelong motor impairments following an early brain injury or malformation (Dan et al. 2025; Baduni et al. 2026). Movement deficits during functional mobility tasks are commonly viewed as a consequence of neuromuscular deficits (Campbell et al. 2024; Licea et al. 2024), but somatosensory impairments are under-recognized contributors to such mobility deficits in CP (Damiano et al. 2013; Santana et al. 2022). Somatosensory dysfunction is a prominent clinical feature in 80–90% of affected individuals (Van Heest et al. 1993; McLean et al. 2021), and a considerable body of work has aimed at determining the neural underpinnings of somatosensory dysfunction in CP (Kurz and Wilson 2011; Maitre et al. 2012; Kurz et al. 2014; Papadelis et al. 2014; Kurz, Becker, et al. 2015; Brun et al. 2021; Knijnenburg et al. 2023; Trevarrow et al. 2023). These investigations have revealed a constellation of atypical neurophysiological features that include attenuated somatosensory cortical responses to transient tactile and electrical stimulation, with weaker responses linked to reduced ankle strength, slower walking speeds and shorter step lengths (Kurz, Heinrichs-Graham, et al. 2015; Trevarrow, Taylor, et al. 2022). Beyond aberrant neural responses to isolated stimuli, people with CP also exhibit altered cortical processing of paired-pulse stimuli, including excessive suppression of the second stimulation (i.e., hypergating) and abnormal secondary cortical responses (Kurz et al. 2018; Trevarrow, Lew, et al. 2022). While such experimental evidence suggests a disruption in the cortical integration of successive sensory inputs, little is known about the capacity of the cortical generators in those with CP to entrain, or neurophysiologically track repetitive sensory stimuli.

Sensory entrainment paradigms provide a rich experimental framework to assess the temporal dynamics of sensory cortical responses that cannot be captured by discrete single- or paired-pulse protocols (Regan 1966; Virlley et al. 2026). These unique paradigms probe the functional properties and integrity of sensory systems using rhythmic, frequency-specific sensory stimulation to capture how neuronal populations synchronize their activity to an exogenous stimulus (Lakatos et al. 2019). The paradigms are generally designed to elicit neural responses that are time- and phased-locked to the stimulus and possess high signal-to-noise ratios, which have driven their widespread use for interrogating both cortical physiology (Heinrichs-Graham and Wilson 2012; Heinrichs-Graham et al. 2017; Wiesman et al. 2019; Wiesman and Wilson 2019; Petro et al. 2024) and neurological dysfunction in clinical populations (Wilson et al. 2007; Wilson et al. 2012; Springer et al. 2022; Chinen et al. 2026; Kurz et al. 2026). These high-fidelity responses are characterized by increases in power (i.e., synchronization) at the fundamental stimulation frequency that evolve over time (Norcia et al. 2015; Heinrichs-Graham *et al*. 2017), reflecting changes in the extent of activation and coordination of the recruited neural generators (Colon et al. 2012; Heinrichs-Graham et al. 2025). Robust entrainment responses are also often observed at the harmonics (i.e., integer multiples of the fundamental frequency) and these may arise from common or distinct neural generators (Kim et al. 2011; Heinrichs-Graham and Wilson 2012) and potentially reflect higher-order nonlinear transformations of sensory inputs (Zhang et al. 2015; Labecki et al. 2016). In addition, concurrent decreases in power (i.e., desynchronization) have also been observed in adjacent spectral bands (Heinrichs-Graham *et al*. 2017), and these are thought to reflect a suppression or shift away from their idling baseline states.

Despite the wealth of neurophysiological insights provided by neural entrainment paradigms, no investigations to date have assessed entrainment in the somatosensory cortices of people with CP. The present study used magnetoencephalography (MEG) to assess neural entrainment to 8 Hz electrical stimulation applied to the posterior tibial nerve in youth with CP and demographically similar neurotypical controls. Given the prior evidence of weaker somatosensory cortical activity in those with CP, we hypothesized that youth with CP would exhibit attenuated and poorly sustained somatosensory entrainment responses compared with the controls. We further aimed to evaluate whether the strength of these cortical responses was related to functional mobility, where we predicted that weaker responses would be related to reduced mobility.

## Materials and Methods

### Participant recruitment

This investigation included youth with spastic CP who had Gross Motor Function Classification System (GMFCS) levels between I-III (Palisano et al. 2008). Individuals at GMFCS level I can walk independently but exhibit difficulty with coordination and balance at faster speeds. Those at GMFCS level II have difficulty walking longer distances and when navigating uneven terrains, and persons at GMFCS level III require mobility aids to walk. Exclusion criteria for the participants with CP were (i) orthopedic surgery within the previous 12 months, (ii) botulinum toxin injections within the previous six months, and (iii) history of epilepsy, seizures, or any psychiatric or neurological disorder unrelated to CP. Clinical phenotypes were determined by licensed physical therapists based on clinical presentation as primary unilateral (UCP) or bilateral (BCP) involvement. Neurotypical controls with no prior psychiatric or neurological disorders, no prior orthopedic surgery, and no ferrous metal implants or MRI safety concerns were similarly recruited. The Institutional Review Board at the Boys Town National Research Hospital approved the study. Prior to study onset, written informed consent was obtained from all participants, and assent was obtained from minors alongside parental consent.

### Clinical assessment of functional mobility

Functional mobility was assessed using the Timed Up and Go (TUG) test that is widely used to index mobility, coordination and dynamic balance in children with CP (Hassani et al. 2014). Participants wore their regular orthotics or shoes and were allowed to use their assistive devices. Seated participants were instructed to stand up, walk at their preferred pace around a cone placed 3 m away, and return to their seated position. Two trials were completed and the fastest completion time was used.

### MEG experimental protocol

Participants sat on a non-magnetic chair in a two-layer magnetically shielded room with their head within a 306-sensor Neo MEG system (MEGIN, Helsinki, Finland) that sampled neuromagnetic signals at 1000 Hz. A constant-current electrical stimulator (Digitimer Limited, Letchworth Garden City, UK) was used to apply a transcutaneous electrical stimulation to the right posterior tibial nerve (pulse width: 200 µs, intensity: 80% of motor threshold). The applied stimulation intensity [median (range)] did not differ between groups [CP = 11 mA (4.40 - 30.0), NT = 9.2 (3.76 - 27.0); Mann Whitney *Z* = - 1.579, *p* = .114]. The stimulation train consisted of pulses applied at 8 Hz for 1000 ms, with an inter-train interval that was randomly jittered between 2600-3400 ms. The 8 Hz entrainment frequency was selected based on pilot work demonstrating that somatosensory cortical entrainment was robust at this frequency (Narici et al. 2001). Participants were instructed to focus on a fixation cross on a screen at eye-level as 100 trials of stimulation were completed.

### MEG acquisition and coregistration with structural MRI

Prior to the MEG experiment, 5 coils were attached to the participant’s head and a 3-dimensional digitizer (Fastrak, Polhemus Navigator Sciences, Colchester, VT, USA) was used to digitize the position of these coils, the three fiducial points, and the scalp surface so that they were within a common head coordinate system, with the location of the coils also being known in reference to the MEG sensor array. This coordinate system was used to coregister the MEG data with the individual’s structural MRI data, which were obtained using a 3T Siemens Prisma scanner equipped with a 64-channel head coil (acquisition parameters: TR = 2400 ms; TE = 1.96 ms; flip angle = 8°; FOV = 256 mm; slice thickness = 1 mm slice with no gap; in-plane resolution = 1.0 mm^3^). Coregistration was performed with the Brain Electrical Source Analysis (BESA) MRI v3.0 software (BESA GmBH, Gräfelfing, Germany). Participant-level MEG data were corrected for head motion and subjected to noise reduction using the signal space separation method with a temporal extension (Taulu and Simola 2006).

### MEG preprocessing, time-frequency transformation and sensor-level statistics

All preprocessing and source imaging of MEG data was conducted using the BESA v7.1 software (BESA GmBH, Gräfelfing, Germany). A bandpass filter (0.5–200 Hz) and notch filter (60 Hz, power mains) was initially applied, and cardiac and eye blink artifacts were removed using the signal-space projection method (Uusitalo and Ilmoniemi 1997). Artifacts induced by the externally applied current were also removed using the extended infomax independent component algorithm (Lee et al. 1999). The resultant continuous neuromagnetic time series data were divided into epochs of 3600 ms length, with stimulation onset at 0 ms, and a baseline period from -1500 to -100 ms. Epochs that contained signal artifacts due to large amplitude or gradient changes were identified using an individualized fixed threshold method supplemented with visual inspection and were rejected from further analysis. The number of retained trials [median (range)] differed across groups (Mann Whitney *Z* = -2.6, *p* = 0.009) with the individuals with CP having fewer retained trials [88 (77–93)] than controls [91.5 (80–97)]. Consequently, the square root of retained trials (SNR) was included as a covariate of no interest in all statistical analyses.

Next, the artifact-free epochs for each sensor were transformed into the time-frequency domain using complex demodulation (0.5 Hz, 100 ms resolution) (Kovach and Gander 2016). The sensor-level data were subsequently averaged across trials and normalized per time-frequency bin using the mean power across the baseline period (-1500 to -100 ms). The specific time-frequency windows of interest were identified based on statistical analyses of the sensor-level spectrograms across the entire array of gradiometers pooled across all participants (Maris and Oostenveld 2007).

### MEG source imaging and virtual sensor analysis

A linearly constrained minimum variance vector beamformer was used to image the time-frequency bins of interest (Gross et al. 2001). By convention, these participant-level source images reflect noise-normalized power differences (*pseudo*-t units, task vs baseline state) per voxel (Hillebrand et al. 2005). These *pseudo*-t functional maps were transformed into standardized Talairach space using the transform that was previously applied to the coregistered structural MRI volume and spatially resampled to 4 mm^3^ resolution. These maps were averaged across all participants to derive grand-averaged beamformer images for the time-frequency bins of interest, and the peak voxels per cluster were used as seeds to extract virtual sensor neural time series data for each participant.

### Statistical analysis

Statistical analyses were conducted in SPSS (v27.0.1, IBM Corp., Armonk, NY) and R Studio (R Core Team, 2025). Normality was assessed using skewness and kurtosis values, and the Shapiro-Wilk test. Group differences in physical characteristics and clinical mobility were assessed using independent t-tests or the non-parametric variant (Mann-Whitney tests), dependent on data normality. Chi-square or Fisher’s exact tests were used for categorical data. Differences in the entrainment temporal dynamics were assessed by extracting relative power values in 100 ms time bins from each virtual sensor time series. As these analyses specifically aimed to capture the temporal evolution of the entrainment neural response, relative power in the first 200 ms was not included. For each time series, separate linear mixed effect models were run with group (CP versus controls) and time (100 ms bins) as fixed factors, and participant as random intercept to account for within-subject correlations arising from repeated measurements. To control for developmental effects and SNR differences between groups, age and SNR were included as covariates of no interest in all models. Age-controlled partial Spearman correlation was used to assess relationships between the mean strength of the cortical responses and the TUG completion time. Of note, additional analyses assessing cross-spectral coupling of the entrainment response at the stimulation frequency and the concurrent alpha response, as well as its association with functional mobility outcomes are described alongside the corresponding results, as their rationale depended on the source imaging outcomes. Significant effects (alpha = 0.05) are reported alongside the magnitude of effect (Cohen’s d (*d*) (Cohen 1988) and were categorized as small (0.2), medium (0.5), or large (≥ 0.8), respectively (Fritz et al. 2012).

## Results

### Participant characteristics

Twenty-eight youth with spastic CP and 34 similar-aged neurotypical controls participated in the study (**Table 1)**. No group differences were observed for age, body mass index, sex, or race distribution (*p*s > 0.05). Overall, youth with CP displayed slower TUG completion times than controls (*d* = 1.17, *p* < .001; **Table 1**), and this was true for those with bilateral (BCP, n = 14; 18.0 ± 10.1 s; *d* = 2.45) and unilateral CP (UCP, n = 14; 6.3 ± 2.0 s; *d* = 1.39; *p*s ≤ .003). Youth with BCP also had slower TUG completion times than those with UCP (*d* = 1.61, *p* = .011).

**Table 1.** Demographic and Physical Characteristics.

| | CP (n = 28) | Controls (n = 34) | $p$ | $d$ |
| --- | --- | --- | --- | --- |
| Age (years) | 21.9 ± 6.3 | 23.4 ± 3.8 | .290 | 0.286 |
| Sex (male/female) | 17/11 | 18/16 | .612 | -- |
| Race<br>(White/Black/Latinx/Asian/Native<br>American/Multiracial) | 25/1/1/1/0/0 | 22/1/2/4/1/4 | .235 | -- |
| BMI (kg/m<sup>2</sup>)* | 24.6 ± 5.8 | 25.6 ± 5.1 | .479 | 0.182 |
| GMFCS level (I/II/III) | 13/6/9 | — | — | — |
| CP presentation<br>(Unilateral/Bilateral) | 14/14 | — | — | — |
| Functional mobility: Timed Up &<br>Go completion time (s) | 12.1 ± 9.3 | <sup>a</sup>4.7 ± 0.5 | <.001 | 1.174 |

*Abbreviations:* BMI, Body mass index; GMFCS, Gross Motor Function Classification System; <sup>a</sup>One participant did not complete the assessment. \*Mann-Whitney $U$ test used.

### Sensor- and source-level results

Sensor level statistical analyses on the spectrograms indicated significant increases in neural response power at the 8 Hz stimulation frequency (i.e., 7.5–8.5 Hz) and its harmonics (second: 15.5–16.5 Hz; third: 23.5–24.5 Hz) that began at stimulation onset (0 ms) and persisted until stimulation offset at 1000 ms (**Fig. 1A**; *p*s < .0001, permutation-corrected). A concurrent decrease in power was also observed within the 10–11 Hz alpha-band from 300 to 1000 ms. For visualization purposes, we also computed group-averaged spectrograms (**Fig. 1B**), which suggested differential patterns of neural responses across the spectrum.

**Figure 1:**
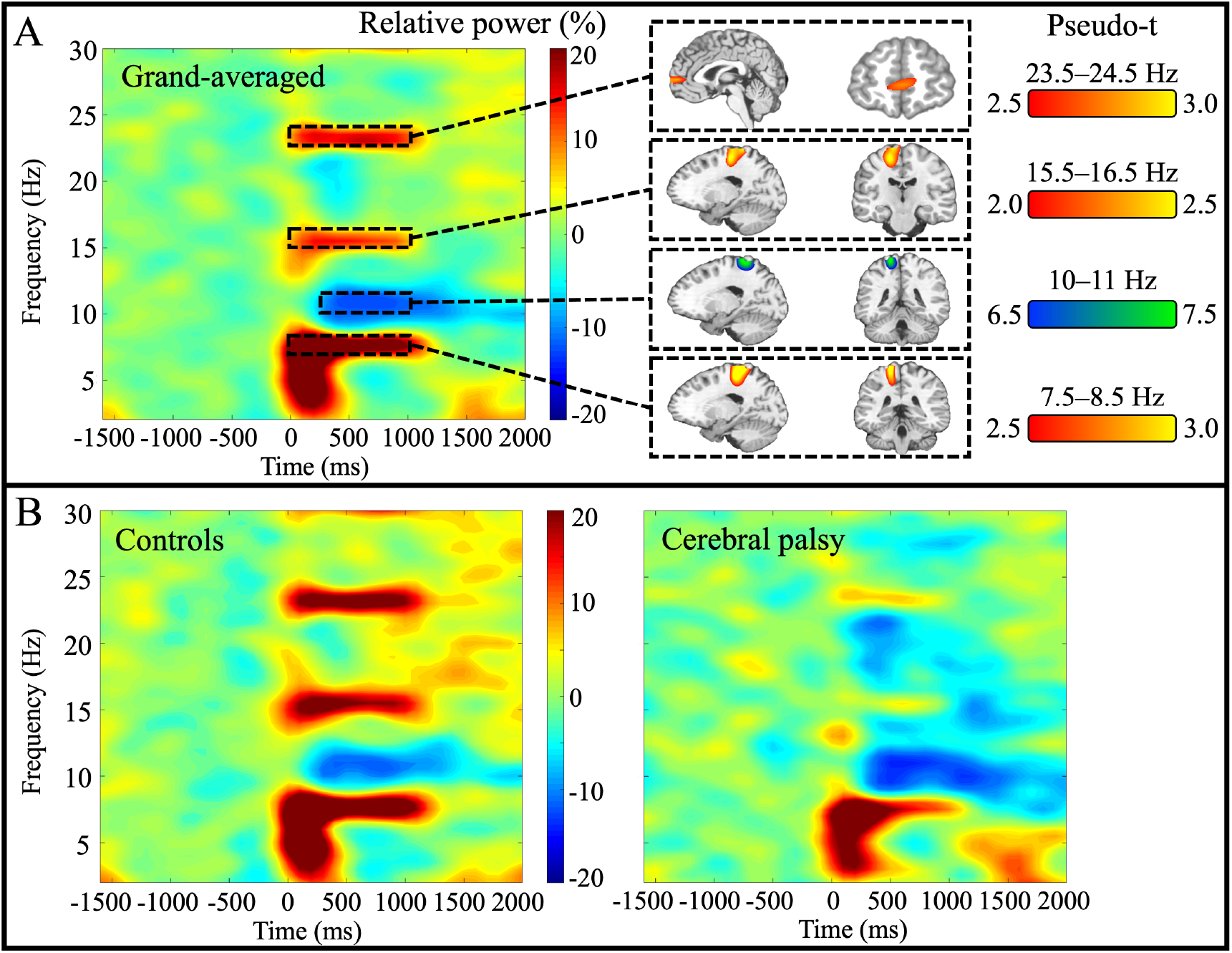
Entrainment responses at the sensor- and source-level. (**A**) The grand-averaged time-frequency spectrogram (n = 62) from MEG sensor 0713, located over the contralateral (i.e., left) sensorimotor cortex, indicated robust increases in power (synchronization) at the entrainment frequency (7.5–8.5 Hz), the second harmonic (15.5–16.5 Hz), and the third harmonic (23.5–24.5 Hz), which spanned the duration of the 8 Hz stimulation (0 to 1000 ms). A concurrent decrease in power (desynchronization) was also observed at the alpha-band (10–11 Hz, 300 to 1000 ms). Beamformer functional maps (in pseudo *t* units, see color bars) revealed the primary sources of these neural responses. The stimulation frequency, second harmonic, and the concurrent alpha response were mapped to the contralateral sensorimotor cortex, while the third harmonic response originated from the prefrontal cortices. (**B**) Group-specific spectrograms from the same sensor for neurotypical controls (n = 34, *left*) and youth with cerebral palsy (n = 28, *right*) are included for visualization purposes, and suggested differential patterns of responses across the spectrum.

Beamformer images of each response were derived for each participant using baselines of equal bandwidth and duration for the synchronization responses (stimulation frequency: 7.5–8.5 Hz, harmonics: 15.5–16.5 and 23.5–24.5 Hz, respectively; all baselines -1200 to -200 ms) and the concurrent alpha response (10–11 Hz, baseline: -1200 to -500 ms). These images were subsequently grand-averaged across participants to identify the brain regions contributing the most to each response. The entrainment responses at the stimulation frequency and second harmonic, and the concurrent alpha response all originated in the contralateral (i.e., left) somatosensory cortex. The entrainment response for the third harmonic was mapped to the bilateral prefrontal cortices.

### Virtual sensor time series analysis

Virtual sensor time series were extracted for each participant using the peak voxel coordinates of the respective grand-averaged source images. The relative power (i.e., change from baseline) time series were then derived in 100 ms time bins from 200–1000 ms for the entrainment responses at the stimulation frequency and both harmonics, and from 300–1000 ms for the concurrent alpha response. These time series were subjected to linear mixed effect modeling with time and group included as fixed factors and participant as random intercept (**Table 2**).

**Table 2.** Summary of linear mixed effects models for relative power changes as a function of group (Cerebral palsy vs controls) and time.

| Parameter | Coefficient ($\beta$) | SE | *t* | *p* |
| --- | --- | --- | --- | --- |
| *Entrainment response:* |  |  |  |  |
| *Stimulation frequency (7.5–8.5 Hz)<sup>a</sup>* |  |  |  |  |
| Group | 3.00 | 4.23 | 0.708 | .481 |
| **Time** | -0.52e<sup>-2</sup> | 0.21e<sup>-2</sup> | -2.52 | **.012** |
| **Group × Time** | -1.29e<sup>-2</sup> | 0.30e<sup>-2</sup> | -4.24 | **<.001** |
| *Concurrent alpha response:* |  |  |  |  |
| *(10–11 Hz)* |  |  |  |  |
| Group | 3.47 | 3.14 | 1.11 | .272 |
| Time | 1.09e<sup>-3</sup> | 1.29e<sup>-3</sup> | 0.84 | .401 |
| **Group × Time** | -8.19e<sup>-3</sup> | 1.93e<sup>-3</sup> | -4.25 | **<.001** |
| *Entrainment response:* |  |  |  |  |
| *Second harmonic (7.5–8.5 Hz)<sup>b</sup>* |  |  |  |  |
| **Group** | -9.93 | 2.57 | -3.86 | **<.001** |
| **Time** | -7.86e<sup>-3</sup> | 1.41e<sup>-3</sup> | -5.55 | **<.001** |
| **Group × Time** | 6.33e<sup>-3</sup> | 2.13e<sup>-3</sup> | 2.98 | **.003** |
| *Entrainment response:* |  |  |  |  |
| *Third harmonic (7.5–8.5 Hz)<sup>c</sup>* |  |  |  |  |
| Group | 0.39 | 0.96 | 0.41 | .683 |
| **Time** | 2.62e<sup>-3</sup> | 9.10e<sup>-4</sup> | 2.88 | **.004** |
| Group × Time | -1.11e<sup>-3</sup> | 9.10e<sup>-4</sup> | -1.22 | 0.223 |

**Bold** font depicts statistically significant group differences at $p < .05$ . All models included mean-centered age and mean-centered SNR (i.e., $\sqrt{\text{accepted trials}}$ ) as fixed factor covariates of no interest, and participant as a random intercept. Outliers identified based on mean power change exceeding $\pm 3$ sd of respective group means: <sup>a</sup> 3 outliers removed, <sup>b</sup> 1 outlier removed, <sup>c</sup> 2 outliers removed.

There was no group main effect for the entrainment response at the stimulation frequency (*p =* .48), indicating that the mean strength of the entrainment response was similar across the two groups (**Fig. 2A**). However, a significant group-by-time interaction (*p* < .001; **Fig. 2B**) indicated a more rapid dissipation of this response over time in the youth with CP compared to the controls.

**Figure 2:**
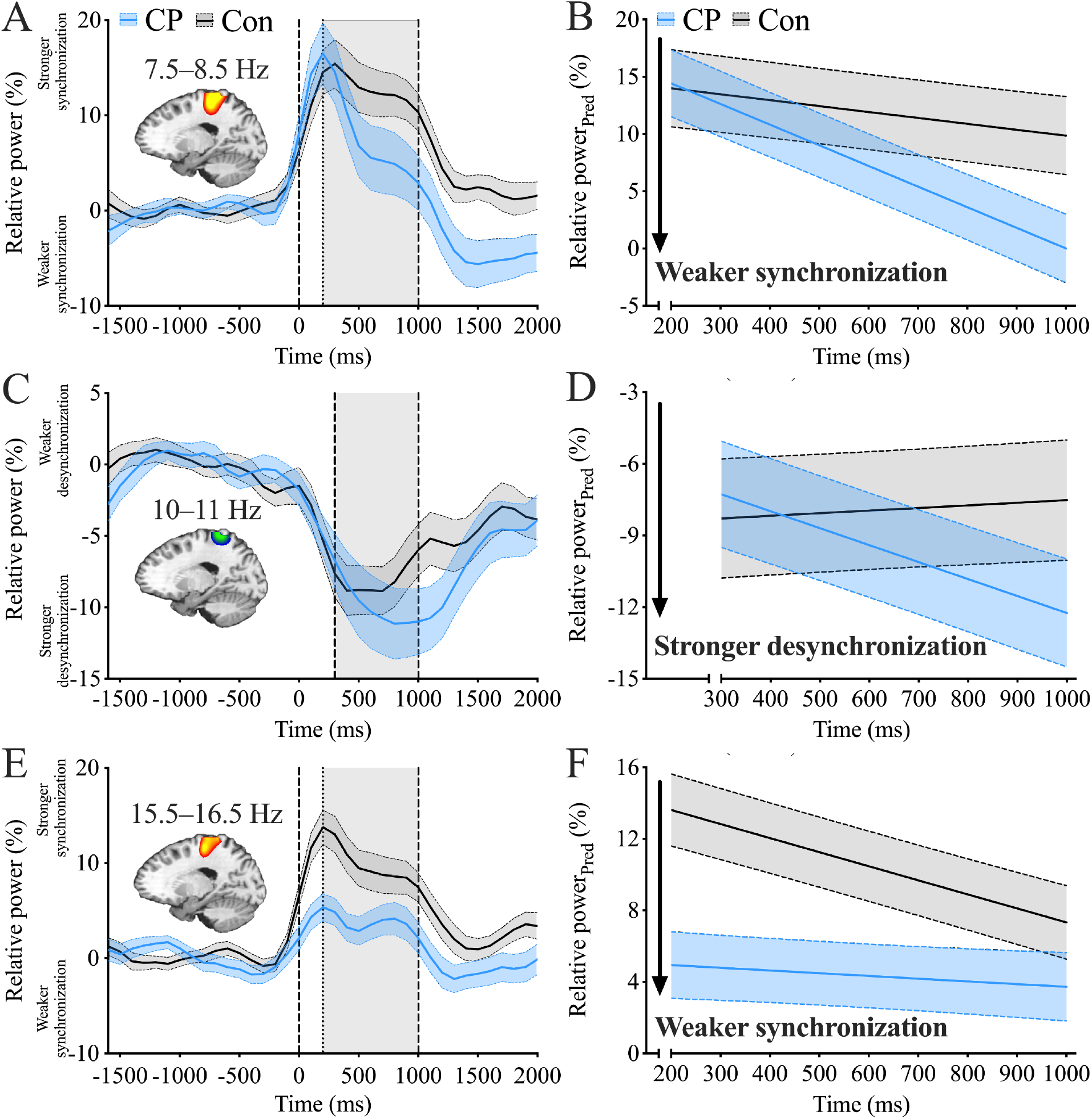
Neural dynamics during somatosensory entrainment. Group-averaged virtual sensor neural time series for relative power (percentage change from baseline units ± SEM in shaded areas) were extracted from the respective peak voxels in the contralateral (i.e., left) somatosensory cortex for the entrainment responses at (**A**) the stimulation frequency (7.5–8.5 Hz) and (**E**) second harmonic band (15.5–16.5 Hz), and (**C**) the concurrent alpha response (10– 11 Hz. Adjacent to each neural time series are group-averaged predicted slopes for change in relative power (estimated marginal means, Relative power_Pred_ on y-axis; mean ± SEM in shaded areas) across specified time windows (dark grey shaded areas in corresponding time series panels, time on x-axis) for **(B)** the entrainment response, **(D)** concurrent alpha response, and (**F**) second harmonic response. Relative to controls, youth with CP exhibited a greater decline in entrainment response, but a greater increase in the concurrent alpha response over time (group-by-time interactions, *p*s < .001). Those with CP also displayed a weaker second harmonic response (group main effect, *p* < .001), alongside a blunted temporal modulation in the strength of this response than that observed in controls (group-by-time interaction, *p* = .003). No significant effects were observed for the third harmonic.

No group effect was observed for the concurrent alpha response (*p =* .27), suggesting that the strength of this response was similar across groups (**Fig. 2C**). However, a significant group-by-time interaction (*p* < .001; **Fig. 2D**) indicated that this response strengthened significantly over time for those with CP, while the controls exhibited a more steady response.

A group main effect was observed for the entrainment response at the second harmonic (*p* < .001), indicating that this response was substantially weaker in those with CP relative to controls (**Fig. 2E**). Additionally, a significant group-by-time interaction (*p* = .003; **Fig. 2F**) indicated that the strength of this response dissipated more sharply in the controls relative to the youth with CP. No group main effect or group-by-time interaction was observed for the entrainment response at the third harmonic (*p*s > .22).

### Cross-spectral coupling of the stimulation frequency entrainment response and the concurrent alpha response

The peak voxels for the respective entrainment response at the stimulation frequency and the concurrent alpha response were spatially adjacent (i.e., in neighboring voxels; Stimulation entrainment: -18, -36, 69; Concurrent alpha: -18, -40, 69), and these responses overlapped in time from 300 to 1000 ms. Therefore, we examined whether the strength of the two responses statistically covaried across the seven overlapping 100 ms time bins. Group differences in this cross-spectral neural coupling were first assessed using an omnibus linear mixed effects model. Concurrent alpha relative power was included as the dependent variable, and entrainment response power, group, time and their interactions as fixed factors, with participant as random intercept, and mean-centered age and SNR as covariates. A significant group-by-entrainment response interaction (*p* = .001) indicated that cross-spectral coupling differed between the groups. Youth with CP displayed a significant positive coupling (*p* < .001) while coupling was not observed in controls (*p* = .636). The group difference in coupling was evident during the later portion of the stimulation (600–1000 ms; *p* = .006) but not the earlier portion (300– 500 ms; *p* = .233), although the group-by-entrainment power-by-time three-way interaction did not reach statistical significance (*p* = .090).

To further characterize the time course of the response coupling, we computed a Pearson correlation coefficient between the two responses pooling across participants within each group at each time bin, followed by a Fisher transformation to enable statistical comparison. These bin-wise comparisons indicated that youth with CP exhibited significant coupling across the 500–1000 ms time bins (one-tailed *t*-tests against zero for Fisher *z* values: *p*s < .030), with significantly higher Fisher *z* (i.e., stronger cross-spectral coupling) than controls across time bins spanning 600–1000 ms (false discovery rate-corrected *p*s ≤ .025; **Fig. 3A**). In contrast, Fisher *z* values in controls did not statistically differ from zero in any time bin (*p*s > .13).

**Figure 3:**
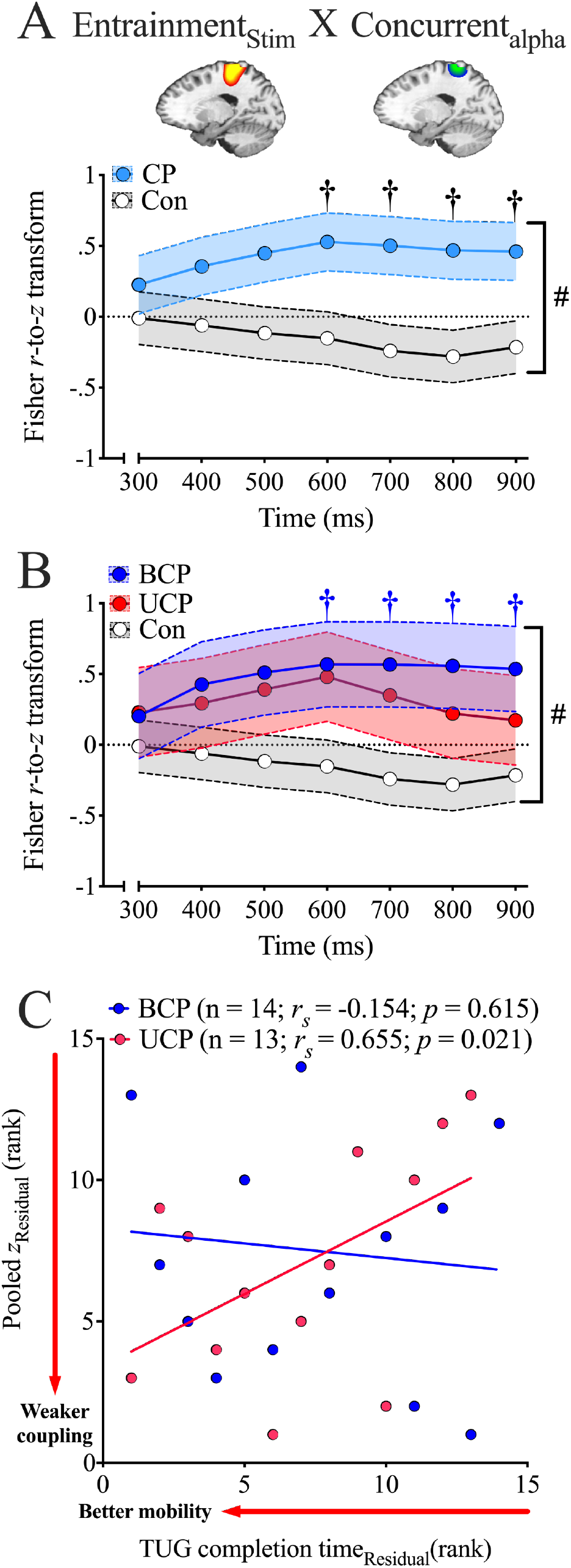
Cross-spectral coupling between the entrainment response at the stimulation frequency (Entrainment_Stim_) and the concurrent alpha response in youth with cerebral palsy (CP) and neurotypical controls (Con), and its association with functional mobility. **(A)** Grand-averaged beamformer images (top) depict the spatially adjacent peak voxels used to seed the virtual sensors for the entrainment response at the stimulation frequency (left, red-yellow) and the concurrent alpha response (right, blue-green) within the contralateral somatosensory cortex. Trial-averaged relative power was extracted from both virtual sensor neural time series across the seven 100 ms time bins. Group-averaged Fisher r-to-z values (mean ± 1 SEM in shaded regions) are plotted per time bin, with positive values indicating a weaker Entrainment_Stim_ response was associated with a stronger concurrent alpha response. Youth with CP (n = 27) exhibited positive Fisher z values from 500 ms onward, whereas values in controls (n = 32) did not differ from zero in any time bin. # denotes a significant group-by-Entrainment_Stim_ response interaction from the omnibus mixed effect model (*p* = .001); † denotes significant group differences at individual time bins (*p*s≤.025, false discovery rate-corrected). (**B**) Exploratory analysis separating the CP group by clinical phenotype. The omnibus group-by-Entrainment_Stim_ interaction was significant (#; *p* < .001), reflecting positive coupling in youth with bilateral CP (BCP; n = 14) that exceeded both unilateral CP (UCP; n = 13; *p* = .011) and controls (*p* < .001). † denotes time bins at which BCP differed from controls (*p*s ≤ .042); Fisher *z* values in UCP did not differ from those of controls or BCP at any bin. (**C**) Overall neural coupling was indexed by a pooled Fisher *z* value computed per participant across all time bins. Age-controlled partial Spearman correlation analysis revealed that youth with UCP who exhibited lower pooled Fisher *z* also tended to display lower TUG completion times (*p* = .021).

We then replicated the preceding omnibus linear mixed effect model-based analyses to explore coupling differences across the CP clinical phenotypes. A significant omnibus group-by-entrainment power interaction (*p* < .001) revealed significant positive coupling specifically in youth with BCP (*p* < .001), which was significantly greater than that in UCP (*p* = .011) and controls (*p* < .001). Further characterization of the time course of this effect per time bin (**Fig. 3B**) indicated that youth with BCP displayed higher Fisher *z* values than controls across the 600–1000 ms time bins (*p*s ≤ .042), but Fisher *z* values in youth with UCP did not differ from those of controls or BCP (*p*s > .05).

### Association of the entrainment neural responses with functional mobility

We first assessed whether the strength of each somatosensory cortical response was associated with functional mobility. Age-controlled partial Spearman revealed no significant associations between the mean strength of neural responses and the TUG completion time for any of the responses in controls or youth with CP (*p*s > .05). We then evaluated if the cross-spectral coupling was associated with functional mobility. A single coefficient of overall cross-spectral neural coupling (pooled Fisher *z*) was derived for each participant by correlating the relative power of their stimulation entrainment and concurrent alpha response across the seven time bins, and applying the same Fisher transformation. Age-controlled partial Spearman correlations indicated that controls who exhibited lower pooled Fisher *z* also tended to display faster TUG completion times (*r_s_* = 0.513; *p* = .004). This relationship was not observed for the entire cohort of participants with CP (*p* = .338), but differed by CP phenotype (**Fig. 3C**). Specifically, youth with UCP who exhibited lower pooled Fisher *z* also tended to display faster TUG completion times (*p* = .021), but this association was not observed in those with BCP (*p* = .615).

## Discussion

The present study provides the first characterization of somatosensory cortical entrainment in youth with CP. Our results indicated that the entrainment response at the stimulation frequency dissipated more quickly in the youth with CP relative to controls, while the concurrent alpha response progressively strengthened over time in CP and remained steady in controls. Cross-spectral coupling between these responses was observed only in youth with CP, was significantly stronger than in controls, and was most prominent in youth with bilateral involvement. Finally, the second harmonic entrainment response was both weaker overall and poorly modulated across time in youth with CP. Collectively, these findings implicate disrupted temporal modulation and atypical cross-spectral coupling of somatosensory cortical oscillatory responses to sustained sensory input as a prominent neurophysiological feature of somatosensory dysfunction in CP.

The first principal finding was the poorly sustained entrainment response at the stimulation frequency for the youth with CP. Both groups rapidly entrained to the stimulation and controls exhibited only mild dissipation in their entrainment response across the duration of the stimulation. Experimental work has shown that repeated sensory stimulation can attenuate cortical responses via short-term depression at thalamocortical synapses (Chung et al. 2002), and cortical entrainment responses can be modeled on thalamocortical connectivity and thalamic activation patterns (Li et al. 2017). In CP, the observed poorly sustained entrainment likely reflects a convergence of multi-level dysfunction across the neuraxis (Coq et al. 2018). Individuals with CP exhibit attenuated somatosensory cortical responses to paired-pulse sensory stimulation, with disproportionate and excessive suppression seen for responses to the secondary pulse (Kurz *et al*. 2018; Trevarrow, Lew*, et al.* 2022). This “hyper-gating” is thought to reflect a compensatory dampening phenomenon, whereby damage to the thalamocortical circuitry (Hoon et al. 2009; Tsao et al. 2015; Papadelis et al. 2019) is thought to contribute to an excitation-inhibition imbalance at the sensory-motor cortices (Illman et al. 2024). Repeated sensory inputs are poorly filtered and inadequately organized at the spinal and thalamic level (Achache et al. 2010; Trevarrow et al. 2021; Dukkipati et al. 2022). Integrating this disorganized signal may plausibly tilt the system toward a cortical suppression bias as a means to reduce computational processing burden. While conceptually plausible, future work is needed to identify whether and how these potential interacting mechanisms contribute to sensory dysfunction in CP.

The second principal finding was that the entrainment response at the stimulation frequency and the concurrent alpha response were significantly coupled in the youth with CP, but did not significantly covary in controls. Exploratory analyses also indicated that the cross-spectral coupling was significantly stronger in youth with BCP than in either UCP or controls. The alpha oscillatory activity represents the dominant idling rhythm of the somatosensory cortex (Salmelin et al. 1995; Hari and Salmelin 1997), and the power reduction is thought to reflect a transition to stimulus-driven active state during somatosensory processing (Cheyne et al. 2003; Pineda 2005; Demas et al. 2020). As such, the entrainment and concurrent alpha responses are presumed to be independently regulated in the intact somatosensory system (Brickwedde et al. 2020). Conversely, the stronger cross-spectral coupling observed in CP may reflect a compensatory phenomenon, whereby individuals with the weakest entrainment may suppress the endogenous alpha rhythm more strongly, indexing a greater engagement of cortical resources when sensory inputs are degraded. Early brain injury in CP may also result in fewer and/or less specialized somatosensory neural ensembles available to sustain the entrainment response, with the loss of specificity or specialization possibly contributing to their degraded function. Alternatively, stronger coupling may also reflect a loss of independent regulation, whereby disrupted subcortical-cortical processing contributes to a functional coupling or yoking together of somatosensory cortical processes that are ordinarily separable in the neurotypical brain. Notably, the present data cannot resolve these speculative accounts, nor can we establish the mechanisms underlying this atypical linkage. Future work integrating novel computational models of the cellular and circuit-level mechanisms of the MEG signals (Jas et al. 2023) may enable testing of these respective viewpoints.

The third principal finding was a weaker entrainment response at the second harmonic in youth with CP, despite preserved overall strength of the entrainment response at the stimulation frequency. Harmonic responses in entrainment paradigms are thought to reflect higher-order nonlinear processing of sensory input (Vlaar et al. 2015; Labecki *et al*. 2016) arising from at least partially distinct set of somatosensory cortical generators (Pastor et al. 2007; Kim *et al*. 2011; Heinrichs-Graham and Wilson 2012; Norcia *et al*. 2015) that may be preferentially attuned to oscillate at beta-band rhythms (Chota et al. 2023). Within this framework, the reduction in CP may reflect weaker engagement of these neural generators by the 8 Hz entrainment stimuli. While the present data cannot empirically discern whether this weaker response reflects fewer neural ensembles entraining to the stimulation, or weaker entrainment of the engaged neural ensembles, the observed patterns broadly align with the attenuated somatosensory cortical responses widely reported in this population (Kurz and Wilson 2011; Kurz *et al*. 2014; Papadelis *et al*. 2014; Kurz, Becker*, et al.* 2015; Kurz *et al*. 2018; Trevarrow, Taylor*, et al.* 2022; Trevarrow *et al*. 2023). A striking parallel is found in entrainment work in the sight-recovery clinical population, wherein individuals with reversed congenital cataracts displayed preserved entrainment responses at the stimulation frequency alongside preferentially attenuated second harmonic responses relative to normal sighted individuals (Pitchaimuthu et al. 2021). This harmonic-specific attenuation was interpreted as preferential involvement of higher-order neural generators in extrastriate cortices that were more sensitive to early developmental deprivation than the primary sensory cortices. The present findings closely follow this pattern, supporting the notion that harmonic generators may be composed of distinct neural ensembles that mediate higher-order sensory processing and are particularly vulnerable to sensorimotor developmental disruption following early brain injury in CP.

Finally, we found that the mean strength of the somatosensory entrainment and concurrent alpha responses was not associated with functional mobility in either CP or controls; however, neural cross-spectral coupling was associated with functional mobility in neurotypical controls and those with UCP. These observations suggest that the temporal organization of these neural responses, rather than their overall magnitude, may better reflect behaviorally relevant somatosensory processing during motor actions. The absence of this relationship in youth with BCP suggests that other factors influenced functional mobility outcomes and likely obscured brain-behavior links in this sub-group. Youth with BCP displayed markedly longer TUG completion times than those with UCP, likely reflecting their more pronounced neuromotor involvement (Damiano et al. 2006) and disproportionate reliance on atypical movement strategies (Goyal et al. 2024). Mobility outcomes in BCP may therefore be driven more by action planning and processing at the motor cortices (Sukal-Moulton et al. 2020) than by somatosensory processing. Given the modest sample size (13–14 participants per subgroup), these CP phenotype-specific associations are preliminary and should at best be considered hypothesis-generating.

### Limitations

The findings of the present study should be interpreted in view of the following limitations. First, while cutaneous electrical stimulation provides a controlled assessment of somatosensory processing, it may not fully capture the complexity of sensory processing during real-world behavior. While overall cross-spectral coupling strength was linked to functional mobility, future work should consider using portable neuroimaging modalities such as electroencephalography (Baker et al. 2020) or optically-pumped magnetometry (OPM-MEG; (Spedden et al. 2025)) to further establish the link between mobility and the altered somatosensory cortical dynamics seen here. Second, the absence of sensory-perceptual measures prevented us from assessing their impact on the study outcomes. Although no gold-standard clinical assessments for somatosensory function currently exist for individuals with CP (Walmsley et al. 2018), future work should consider including clinical sensory-perceptual evaluations for improved generalizability. Third, the sub-group analyses across the phenotypes of CP were exploratory and likely underpowered to detect small-to-moderate brain–behavior associations. As such, the absence of a neural coupling–mobility relationship in youth with BCP may be interpreted as hypothesis-generating. Finally, phenotypes of CP were categorized on clinical presentation rather than brain injury patterns. This approach was intentional, as structural injury patterns are variably linked with clinical presentation and functional outcomes in CP (Pagnozzi et al. 2020). Future work should aim to delineate the underlying mechanisms of the present findings.

### Conclusion

This study is the first to characterize somatosensory cortical entrainment in individuals with CP, and reveals disrupted temporal dynamics of cortical responses to repeated posterior tibial nerve stimulation. Youth with CP displayed poorly sustained entrainment at the stimulation frequency, as well as atypical concurrent alpha responses that progressively strengthened rather than stabilizing over time. Importantly, these two responses were statistically coupled in youth with CP but remained dissociable in controls. This cross-spectral coupling was linked to functional mobility in controls and, in exploratory analyses, appeared confined to youth with unilateral CP. Finally, the entrainment responses at the second harmonic were markedly attenuated in CP, suggesting preferential susceptibility of these higher-order neural generators to disruption from early brain injury. Collectively, these observations reveal aberrant somatosensory cortical entrainment responses following early brain injury in youth with CP, and motivate further work to determine whether these neurophysiological features are modifiable with rehabilitative interventions.

## Author contributions statement

**O.A.K.**: Writing – original draft, Writing – review & editing, Investigation,Formal analysis, Visualization, Validation. **S.E.B.**: Writing – review & editing, Data curation, Investigation, Validation. **E.H-G.**: Writing – review & editing, Methodology. **T.W.W.**: Writing – review & editing, Software, Resources, Funding acquisition. **M.J.K.**: Writing – review & editing, Supervision, Resources, Methodology, Funding acquisition, Conceptualization, Project administration.

## Data availability

Data will be made available upon reasonable request.

## References

Achache V, Roche N, Lamy JC, Boakye M, Lackmy A et al. 2010. Transmission within several spinal pathways in adults with cerebral palsy. Brain. 133:1470–1483.

Baduni K, Khan OA, Modlesky CM, Maitre NL. 2026. Early motor and cognitive development in typically developing children and those with or at high risk of cerebral palsy: A scoping review. Developmental medicine and child neurology. 68:623–635.

Baker S, Trevarrow M, Gehringer J, Bergwell H, Arpin D et al. 2020. Gamma somatosensory cortical oscillations are attenuated during the stance phase of human walking. Neurosci Lett. 732:135090.

Brickwedde M, Schmidt MD, Kruger MC, Dinse HR. 2020. 20 Hz Steady-State Response in Somatosensory Cortex During Induction of Tactile Perceptual Learning Through LTP-Like Sensory Stimulation. Front Hum Neurosci. 14:257.

Brun C, Traverse E, Granger E, Mercier C. 2021. Somatosensory deficits and neural correlates in cerebral palsy: a scoping review. Developmental medicine and child neurology. 63:1382–1393.

Campbell KS, Whitten SVW, Newell KM, Li L, Singh T et al. 2024. Deficits in recovery of postural stability after stepping are limb- and phase-specific in children with unilateral cerebral palsy. Gait & Posture. 114:227–233.

Cheyne D, Gaetz W, Garnero L, Lachaux JP, Ducorps A et al. 2003. Neuromagnetic imaging of cortical oscillations accompanying tactile stimulation. Brain Res Cogn Brain Res. 17:599–611.

Chinen L, Busboom MT, Meejang J, Kastner O, Heinrichs-Graham E et al. 2026. Adults with Down syndrome display altered entrainment of occipital cortical neurons. Brain Commun. 8:fcag038.

Chota S, VanRullen R, Gulbinaite R. 2023. Random Tactile Noise Stimulation Reveals Beta-Rhythmic Impulse Response Function of the Somatosensory System. J Neurosci. 43:3107–3119.

Chung S, Li X, Nelson SB. 2002. Short-term depression at thalamocortical synapses contributes to rapid adaptation of cortical sensory responses in vivo. Neuron. 34:437–446.

Cohen J. 1988. Statistical power analysis for the behavioral sciences. Hillsdale, N.J.: Routledge.

Colon E, Legrain V, Mouraux A. 2012. Steady-state evoked potentials to study the processing of tactile and nociceptive somatosensory input in the human brain. Neurophysiol Clin. 42:315–323.

Coq JO, Delcour M, Ogawa Y, Peyronnet J, Castets F et al. 2018. Mild Intrauterine Hypoperfusion Leads to Lumbar and Cortical Hyperexcitability, Spasticity, and Muscle Dysfunctions in Rats: Implications for Prematurity. Front Neurol. 9:423.

Damiano D, Abel M, Romness M, Oeffinger D, Tylkowski C et al. 2006. Comparing functional profiles of children with hemiplegic and diplegic cerebral palsy in GMFCS Levels I and II: Are separate classifications needed? Developmental medicine and child neurology. 48:797–803.

Damiano DL, Wingert JR, Stanley CJ, Curatalo L. 2013. Contribution of hip joint proprioception to static and dynamic balance in cerebral palsy: a case control study. J Neuroeng Rehabil. 10:57.

Dan B, Rosenbaum P, Carr L, Gough M, Coughlan J et al. 2025. Proposed updated description of cerebral palsy. Developmental medicine and child neurology. 67:700–709.

Demas J, Bourguignon M, Perivier M, De Tiege X, Dinomais M et al. 2020. Mu rhythm: State of the art with special focus on cerebral palsy. Ann Phys Rehabil Med. 63:439–446.

Dukkipati SS, Walker SJ, Trevarrow MP, Busboom M, Baker SE et al. 2022. Reduced wrist flexor H-reflex excitability is linked with increased wrist proprioceptive error in adults with cerebral palsy. Front Neurol. 13:930303.

Fritz CO, Morris PE, Richler JJ. 2012. Effect size estimates: current use, calculations, and interpretation. J Exp Psychol Gen. 141:2–18.

Goyal V, Gordon KE, Sukal-Moulton T. 2024. Children with bilateral cerebral palsy use their hip joint to complete a step-up task. Front Hum Neurosci. 18:1343457.

Gross J, Kujala J, Hamalainen M, Timmermann L, Schnitzler A et al. 2001. Dynamic imaging of coherent sources: Studying neural interactions in the human brain. Proceedings of the National Academy of Sciences. 98:694–699.

Hari R, Salmelin R. 1997. Human cortical oscillations: a neuromagnetic view through the skull. Trends Neurosci. 20:44–49.

Hassani S, Krzak JJ, Johnson B, Flanagan A, Gorton G, 3rd et al. 2014. One-Minute Walk and modified Timed Up and Go tests in children with cerebral palsy: performance and minimum clinically important differences. Developmental medicine and child neurology. 56:482–489.

Heinrichs-Graham E, McDermott TJ, Mills MS, Coolidge NM, Wilson TW. 2017. Transcranial direct-current stimulation modulates offline visual oscillatory activity: A magnetoencephalography study. Cortex. 88:19–31.

Heinrichs-Graham E, Shen Z, Lee WH, Benavente AA, Eastman JA et al. 2025. Cross-modal impact of auditory experience on visual cortical entrainment in children who are hard-of-hearing. Imaging Neurosci (Camb). 3.

Heinrichs-Graham E, Wilson TW. 2012. Presence of strong harmonics during visual entrainment: a magnetoencephalography study. Biol Psychol. 91:59–64.

Hillebrand A, Singh KD, Holliday IE, Furlong PL, Barnes GR. 2005. A new approach to neuroimaging with magnetoencephalography. Hum Brain Mapp. 25:199–211.

Hoon, AH Jr., Stashinko EE, Nagae LM, Lin DD, Keller J et al. 2009. Sensory and motor deficits in children with cerebral palsy born preterm correlate with diffusion tensor imaging abnormalities in thalamocortical pathways. Developmental medicine and child neurology. 51:697–704.

Illman M, Jaatela J, Vallinoja J, Nurmi T, Maenpaa H et al. 2024. Altered excitation-inhibition balance in the primary sensorimotor cortex to proprioceptive hand stimulation in cerebral palsy. Clin Neurophysiol. 157:25–36.

Jas M, Thorpe R, Tolley N, Bailey C, Brandt S et al. 2023. HNN-core: A Python software for cellular and circuit-level interpretation of human MEG/EEG. J Open Source Softw. 8.

Kim YJ, Grabowecky M, Paller KA, Suzuki S. 2011. Differential roles of frequency-following and frequency-doubling visual responses revealed by evoked neural harmonics. J Cogn Neurosci. 23:1875–1886.

Knijnenburg ACS, Steinbusch CVM, Janssen-Potten YJM, Defesche A, Vermeulen RJ. 2023. Neuro-imaging characteristics of sensory impairment in cerebral palsy; a systematic review. Front Rehabil Sci. 4:1084746.

Kovach CK, Gander PE. 2016. The demodulated band transform. Journal of Neuroscience Methods. 261:135–154.

Kurz MJ, Becker KM, Heinrichs-Graham E, Wilson TW. 2015. Children with cerebral palsy have uncharacteristic somatosensory cortical oscillations after stimulation of the hand mechanoreceptors. Neuroscience. 305:67–75.

Kurz MJ, Dao E, Busboom MT, Heinrichs-Graham E, Corr B et al. 2026. Persons with cerebral palsy display improved occipital cortical entrainment after gait training. Neuroscience. 595:73–80.

Kurz MJ, Heinrichs-Graham E, Arpin DJ, Becker KM, Wilson TW. 2014. Aberrant synchrony in the somatosensory cortices predicts motor performance errors in children with cerebral palsy. J Neurophysiol. 111:573–579.

Kurz MJ, Heinrichs-Graham E, Becker KM, Wilson TW. 2015. The magnitude of the somatosensory cortical activity is related to the mobility and strength impairments seen in children with cerebral palsy. J Neurophysiol. 113:3143–3150.

Kurz MJ, Wiesman AI, Coolidge NM, Wilson TW. 2018. Children with Cerebral Palsy Hyper-Gate Somatosensory Stimulations of the Foot. Cereb Cortex. 28:2431–2438.

Kurz MJ, Wilson TW. 2011. Neuromagnetic activity in the somatosensory cortices of children with cerebral palsy. Neurosci Lett. 490:1–5.

Labecki M, Kus R, Brzozowska A, Stacewicz T, Bhattacharya BS et al. 2016. Nonlinear Origin of SSVEP Spectra-A Combined Experimental and Modeling Study. Front Comput Neurosci. 10:129.

Lakatos P, Gross J, Thut G. 2019. A New Unifying Account of the Roles of Neuronal Entrainment. Curr Biol. 29:R890–R905.

Lee TW, Girolami M, Sejnowski TJ. 1999. Independent component analysis using an extended infomax algorithm for mixed subgaussian and supergaussian sources. Neural Comput. 11:417–441.

Li G, Henriquez CS, Frohlich F. 2017. Unified thalamic model generates multiple distinct oscillations with state-dependent entrainment by stimulation. PLoS Comput Biol. 13:e1005797.

Licea J, Khan OA, Singh T, Modlesky CM. 2024. Prefrontal cortex hemodynamic activity during a test of lower extremity functional muscle strength in children with cerebral palsy: A functional near-infrared spectroscopy study. Eur J Neurosci. 59:298–307.

Maitre NL, Barnett ZP, Key AP. 2012. Novel assessment of cortical response to somatosensory stimuli in children with hemiparetic cerebral palsy. J Child Neurol. 27:1276–1283.

Maris E, Oostenveld R. 2007. Nonparametric statistical testing of EEG- and MEG-data. Journal of Neuroscience Methods. 164:177–190.

McLean B, Taylor S, Valentine J, Carey L, Thornton A et al. 2021. Somatosensory discrimination impairment in children with hemiplegic cerebral palsy as measured by the sense_assess(c) kids. Aust Occup Ther J. 68:317–326.

Narici L, Forss N, Jousmaki V, Peresson M, Hari R. 2001. Evidence for a 7- to 9-Hz “sigma” rhythm in the human SII cortex. Neuroimage. 13:662–668.

Norcia AM, Appelbaum LG, Ales JM, Cottereau BR, Rossion B. 2015. The steady-state visual evoked potential in vision research: A review. J Vis. 15:4.

Pagnozzi AM, Pannek K, Fripp J, Fiori S, Boyd RN et al. 2020. Understanding the impact of bilateral brain injury in children with unilateral cerebral palsy. Hum Brain Mapp. 41:2794–2807.

Palisano RJ, Rosenbaum P, Bartlett D, Livingston MH. 2008. Content validity of the expanded and revised Gross Motor Function Classification System. Developmental medicine and child neurology. 50:744–750.

Papadelis C, Ahtam B, Feldman HA, AlHilani M, Tamilia E et al. 2019. Altered White Matter Connectivity Associated with Intergyral Brain Disorganization in Hemiplegic Cerebral Palsy. Neuroscience. 399:146–160.

Papadelis C, Ahtam B, Nazarova M, Nimec D, Snyder B et al. 2014. Cortical somatosensory reorganization in children with spastic cerebral palsy: a multimodal neuroimaging study. Front Hum Neurosci. 8:725.

Pastor MA, Valencia M, Artieda J, Alegre M, Masdeu JC. 2007. Topography of cortical activation differs for fundamental and harmonic frequencies of the steady-state visual-evoked responses. An EEG and PET H215O study. Cereb Cortex. 17:1899–1905.

Petro NM, Webert LK, Springer SD, Okelberry HJ, John JA et al. 2024. Optimal gamma-band entrainment of visual cortex. Hum Brain Mapp. 45:e26775.

Pineda JA. 2005. The functional significance of mu rhythms: translating “seeing” and “hearing” into “doing”. Brain Res Brain Res Rev. 50:57–68.

Pitchaimuthu K, Dormal G, Sourav S, Shareef I, Rajendran SS et al. 2021. Steady state evoked potentials indicate changes in nonlinear neural mechanisms of vision in sight recovery individuals. Cortex. 144:15–28.

Regan D. 1966. Some characteristics of average steady-state and transient responses evoked by modulated light. Electroencephalogr Clin Neurophysiol. 20:238–248.

Salmelin R, Hamalainen M, Kajola M, Hari R. 1995. Functional segregation of movement-related rhythmic activity in the human brain. Neuroimage. 2:237–243.

Santana CAS, Dos Santos MM, de Campos AC. 2022. Interrelationships of Touch and Proprioception with Motor Impairments in Individuals with Cerebral Palsy: A Systematic Review. Percept Mot Skills. 129:570–590.

Spedden ME, O’Neill GC, West TO, Tierney TM, Mellor S et al. 2025. Using Wearable MEG to Study the Neural Control of Human Stepping. Sensors (Basel). 25.

Springer SD, Wiesman AI, May PE, Schantell M, Johnson HJ et al. 2022. Altered visual entrainment in patients with Alzheimer’s disease: magnetoencephalography evidence. Brain Commun. 4:fcac198.

Sukal-Moulton T, de Campos AC, Alter KE, Damiano DL. 2020. Functional near-infrared spectroscopy to assess sensorimotor cortical activity during hand squeezing and ankle dorsiflexion in individuals with and without bilateral and unilateral cerebral palsy. Neurophoton. 7:045001.

Taulu S, Simola J. 2006. Spatiotemporal signal space separation method for rejecting nearby interference in MEG measurements. Phys Med Biol. 51:1759–1768.

Trevarrow MP, Baker SE, Wilson TW, Kurz MJ. 2021. Microstructural changes in the spinal cord of adults with cerebral palsy. Developmental medicine and child neurology. 63:998–1003.

Trevarrow MP, Bergwell HE, Groff BR, Wiesman AI, Wilson TW et al. 2023. Youth with Cerebral Palsy Display Abnormal Somatosensory Cortical Activity During a Haptic Exploration Task. Neuroscience. 515:53–61.

Trevarrow MP, Lew BJ, Hoffman RM, Taylor BK, Wilson TW et al. 2022. Altered Somatosensory Cortical Activity Is Associated with Cortical Thickness in Adults with Cerebral Palsy: Multimodal Evidence from MEG/sMRI. Cereb Cortex. 32:1286–1294.

Trevarrow MP, Taylor BK, Reelfs AM, Wilson TW, Kurz MJ. 2022. Aberrant movement-related somatosensory cortical activity mediates the extent of the mobility impairments in persons with cerebral palsy. J Physiol. 600:3537–3548.

Tsao H, Pannek K, Boyd RN, Rose SE. 2015. Changes in the integrity of thalamocortical connections are associated with sensorimotor deficits in children with congenital hemiplegia. Brain Struct Funct. 220:307–318.

Uusitalo MA, Ilmoniemi RJ. 1997. Signal-space projection method for separating MEG or EEG into components. Med Biol Eng Comput. 35:135–140.

Van Heest AE, House J, Putnam M. 1993. Sensibility deficiencies in the hands of children with spastic hemiplegia. J Hand Surg Am. 18:278–281.

Virlley M, Spooner RK, Wilson TW, Pruitt T, Davenport EM et al. 2026. Gating of somatosensory neural oscillations across the lifespan: A narrative review. Ageing Res Rev. 121:103276.

Vlaar MP, van der Helm FCT, Schouten AC. 2015. Frequency Domain Characterization of the Somatosensory Steady State Response in Electroencephalography. IFAC-PapersOnLine. 48:1391–1396.

Walmsley C, Taylor S, Parkins T, Carey L, Girdler S et al. 2018. What is the current practice of therapists in the measurement of somatosensation in children with cerebral palsy and other neurological disorders? Aust Occup Ther J. 65:89–97.

Wiesman AI, Groff BR, Wilson TW. 2019. Frontoparietal Networks Mediate the Behavioral Impact of Alpha Inhibition in Visual Cortex. Cereb Cortex. 29:3505–3513.

Wiesman AI, Wilson TW. 2019. Alpha Frequency Entrainment Reduces the Effect of Visual Distractors. J Cogn Neurosci. 31:1392–1403.

Wilson TW, Rojas DC, Reite ML, Teale PD, Rogers SJ. 2007. Children and adolescents with autism exhibit reduced MEG steady-state gamma responses. Biol Psychiatry. 62:192–197.

Wilson TW, Wetzel MW, White ML, Knott NL. 2012. Gamma-frequency neuronal activity is diminished in adults with attention-deficit/hyperactivity disorder: a pharmaco-MEG study. J Psychopharmacol. 26:771–777.

Zhang Y, Guo D, Cheng K, Yao D, Xu P. 2015. The graph theoretical analysis of the SSVEP harmonic response networks. Cogn Neurodyn. 9:305–315.

